# Enhancing Drug Delivery with Supramolecular Amphiphilic Macrocycle Nanoparticles: Selective Targeting of CDK4/6 Inhibitor Palbociclib to Melanoma

**DOI:** 10.1101/2023.11.21.567974

**Authors:** Mohamed F. Attia, Edikan A. Ogunnaike, Megan Pitz, Nancy M. Elbaz, Dillip K. Panda, Angela Alexander-Bryant, Sourav Saha, Daniel C. Whitehead, Alexander Kabanov

**Affiliations:** Center for Nanotechnology in Drug Delivery and Division of Pharmacoengineering and Molecular Pharmaceutics, Eshelman School of Pharmacy, University of North Carolina at Chapel Hill, NC 27599, USA; Department of Bioengineering, Clemson University, Clemson, SC, 29634, USA; Department of Chemistry, Clemson University, Clemson, SC, 29634, USA; Laboratory of Chemical Design of Bionanomaterials, Faculty of Chemistry, M.V. Lomonosov Moscow State University, Moscow, 119992, Russia

**Keywords:** Supramolecular macrocycles, nanoparticle drug delivery, CDK4/6 inhibitor, Palbociclib, melanoma, glioblastoma, cancer

## Abstract

Drug delivery systems based on amphiphilic supramolecular macrocycles have garnered increased attention over the past two decades due to their ability to successfully formulate nanoparticles. Macrocyclic (MC) materials can self-assemble at lower concentrations without the need for surfactants and polymers, but surfactants are required to form and stabilize nanoparticles at higher concentrations. Using MCs to deliver both hydrophilic and hydrophobic guest molecules is advantageous. We developed two novel types of amphiphilic macrocycle nanoparticles (MC NPs) capable of delivering either Nile Red (NR) (a hydrophobic model) or Rhodamine B (RhB) (a hydrophilic model) fluorescent dyes. We extensively characterized the materials using various techniques to determine size, morphology, stability, hemolysis, fluorescence, loading efficiency (LE), and loading capacity (LC). We then loaded the CDK4/6 inhibitor Palbociclib (Palb) into both MC NPs using a solvent diffusion method. This yielded Palb-MC NPs in the size range of 65-90 nm. They exhibited high stability over time and in fetal bovine serum with negligible toxicity against erythrocytes. Cytotoxicity was minimal when tested against RAW macrophages, <u>human fibroblast HDFn</u>, and adipose stromal cells (ASCs) at higher concentrations of MC NPs. Cell viability studies were conducted with different concentrations of MC NPs, Palb-MC NPs, and free Palb against RAW macrophages, human U-87 GBM, and human M14 melanoma cell lines in vitro. Flow cytometry experiments revealed that blank MC NPs and Palb-MC NPs were selectively targeted to melanoma cells, resulting in cell death compared to the other two cell lines. Future work will focus on studying the biological effect of MC NPs including their binding affinity with molecules/receptors expressed on the M14 and other melanoma cell surface by molecular docking simulations. Subsequently, we will evaluate the MCs as a component of combination therapy in a murine melanoma model.

**Graphical abstract:** 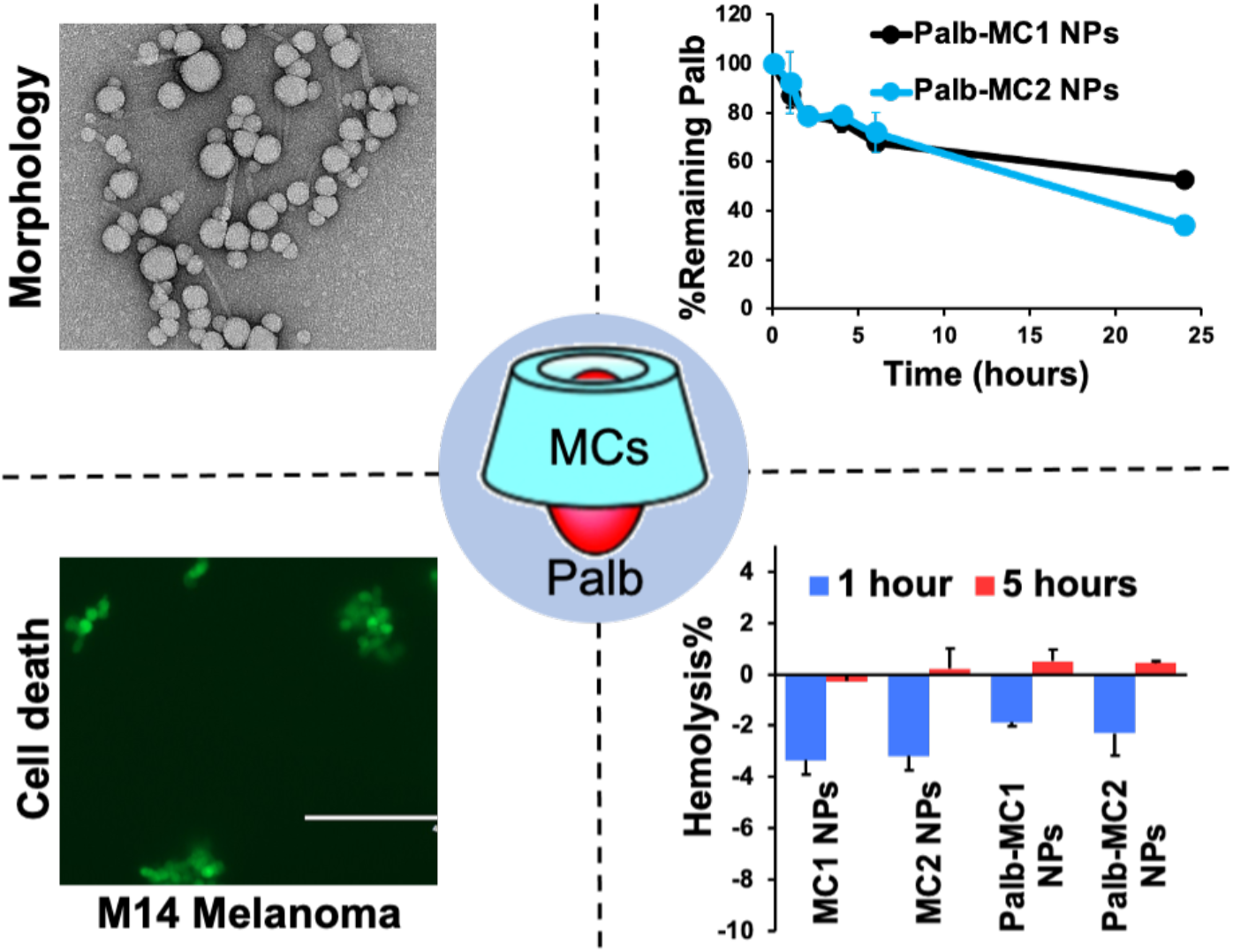

## 1. Introduction

Our goal in this study was to evaluate novel (MC) compounds as a potential drug delivery system for inhibitors of cyclin-dependent kinases (CDK) to treat metastatic melanoma. Metastatic melanoma is known to be a highly fatal cancer,^1^ but the advent of more effective therapies including BRAF/MEK inhibitors and anti-PD-1 antibodies is beginning to turn the tide. Nonetheless, some patients still fail to respond to these therapies. The dysregulation of cell cycle progression is correlated to tumorigenesis, and compounds that prevent cell proliferation by inhibiting CDK have been recognized as an effective anticancer therapy.^2^

The use of supramolecular MC host-guest complexes is an attractive strategy for drug encapsulation, as it can improve solubility and enable controlled drug release.^3, 4^ This approach relies on noncovalent interactions, typically hydrogen bonding, in aqueous environments to generate supramolecular complexes that encapsulate hydrophobic drugs within the macrocycle cavity. In turn, the macrocycle’s outer shell exposes hydrophilic sites.^5^ Previous studies have demonstrated that macrocyclic systems can be customized to incorporate multiple drug moieties and display targeting elements on the construct’s surface.^6, 7^

The MC amphiphiles utilized in our study were first synthesized by Mitra et al.^8^, who described their self-assembly into various supramolecular nanostructures. These MC amphiphiles are unique materials capable of loading both hydrophilic and hydrophobic guest molecules. Their uniqueness lies in their ability to form stable host-guest complexes through hydrophobic interactions and/or hydrogen bonding between the host (MCs) and the guest (small drugs). Additionally, MCs can form three-dimensional and porous networks or multiphase structures, which facilitate the chemical and/or physical entrapment and controlled release of therapeutic agents. Another advantage of these multifunctional materials is their ability to conjugate targeting moieties to MC amphiphiles for site-specific delivery of guest molecules. Furthermore, MCs can exhibit pharmacological effects that potentially enhance the effects of drug/gene delivery. However, the pharmacological evaluation of these materials for drug delivery has not been reported to the best of our knowledge.

In this study, we report the successful formulation of stable nanoparticles using our novel synthesized amphiphilic supramolecular MCs containing nitro groups (MC1) or amino groups (MC2). These MC NPs were easily prepared using the solvent diffusion method with the assistance of a polyethoxylated nonionic surfactant as a solubilizer. The MC NPs displayed the ability to load both hydrophilic and hydrophobic fluorescent dyes, which were used as model drugs. Building on these initial positive results, we further investigated the potential of macrocyclic (MC) compounds to enhance the delivery of Palbociclib (Palb), a CDK4/6 inhibitor that controls the progression of the cell cycle. CDK4/6 activation induces the phosphorylation of retinoblastoma protein (pRb), leading to its inactivation and preventing cell cycle progression from the G1/1 phase to the S phase.^9, 10^ This makes them potential targets in treating various cancers, including breast cancer, melanoma, and glioblastoma multiforme (GBM).^11–14^ Palb is a targeted drug therapy known for causing fewer unwanted side effects compared to traditional chemotherapy.

Our results showed that Palb-loaded MC NPs exhibited improved biocompatibility, stability, and a slow drug release profile, which enhanced the utility of Palb. We evaluated the performance of Palb-loaded MCs on three cell lines: RAW 264.7 macrophages, U-87 GBM, and M14 melanoma cells. Our results demonstrate that both blank MC NPs and Palb-MC NPs displayed enhanced cytotoxicity towards melanoma cells and induced significant cell death *in vitro* compared to the other two cell lines. These findings are promising and provide valuable insights into the potential of MC NPs as a drug delivery system for the treatment of melanoma tumors. Furthermore, these results open up avenues for further investigation and evaluation of the efficacy of Palb-MC NPs in a murine melanoma model.

## 2. Materials and Methods

### Chemicals and reagents

<u>Palb-free</u> base and Palb-HCl salt were obtained from LC Laboratories (Woburn, MA). Nile red (NR) and rhodamine B (RhB) dyes, dimethyl sulfoxide (DMSO), and tetrahydrofuran (THF) were purchased from Sigma-Aldrich (St. Louis, MO). A non-ionic PEGylated surfactant (Kolliphor ELP^®^ formerly known as Cremophor EL (CrEL), which is a polyethoxylated castor oil derivative was purchased from BASF (Ludwigshafen, Germany). Water and acetonitrile (HPLC grade) were purchased from Fisher Scientific Inc. Phosphate-buffered saline (PBS), Dulbecco’s modified Eagle’s medium (DMEM), Roswell Park Memorial Institute (RPMI) 1640 Medium, fetal bovine serum (FBS), penicillin/streptomycin, <u>CellMask Plasma Membrane stain</u>, and Cell Counting Kit-8 (CCK-8) were purchased from Fisher Scientific. Syringe filters (PTFE 0.4 µm), centrifugal units (10 kDa cut off), and 3.5 kDa floatable slide-A-lyzer Mini dialysis devices were purchased from Thermo Fisher Scientific.

### Cell lines and cell culture

We obtained GFP-expressing cells from Gianpietro Dotti’s laboratory (Lineberger Comprehensive Cancer Center, Chapel Hill, NC, USA) [Research Resource Identifiers (#RRID):CVCL 0022]. Specifically, we obtained human melanoma M14 cells (source: JWCI) expressing GFP (M14-GFP) and the GBM tumor cell line U-87 MG (source: male) expressing GFP (U-87 GBM-GFP). We also obtained RAW 264.7 mouse macrophages, U-87 MG, and adipose stromal cells, <u>human dermal fibroblasts, neonatal (HDFn), human melanoma (Mel29.1) cells</u> from the American Type Culture Collection (ATCC). We purchased Invitrogen NucBlue live cell nuclear stain from Thermo Fisher Scientific. The cells were grown in RPMI 1640 or DMEM supplemented with 10% FBS (Gemini Bio-Products), 1% penicillin/streptomycin, and 1% GlutaMAX. They were maintained in a humidified atmosphere containing 5% CO_2_ at 37 °C.

### Synthesis of supramolecular macrocycles MC1 and MC2

The synthesis and characterization of MC1 and MC2 amphiphilic macrocycles have been previously described.^8^

### Preparation of MC1 and MC2 NPs

Supramolecular macrocyclic NPs were prepared by solvent diffusion method as described elsewhere.^15^ Briefly, a stock solution of organic phase containing the MC1 or MC2 material (2 mg) and CrEL (3.6 mg) in THF or DMSO (200 µL) were mixed together by vortex, heating, and sonication until complete homogenization. This organic phase was dropwise added to 1800 µL of deionized (DI) water and kept stirring at 700 rpm for one hour at room temperature. Next, a purification process was done to remove the free molecules and organic solvent from the solution by using a syringe filter (0.45 µm) followed by centrifugation at 3500 rpm for 20 min using centrifugal filter units (10 kDa cutoff). The washing process is repeated three times using DI water and concentrates the final solution to 200 µL making the final concentration of MC material 10 mg/mL. Preparation of MC1 or MC2 nanoparticles loaded with cargo was also done by insertion of different concentrations of guest molecules (*i.e.* fluorescent NR, RhB, or Palb) within the organic phase following previous steps.

### Dynamic Light Scattering (DLS)

The sizes of both free and cargo-loaded MC particles, as well as their polydispersity index (PDI) and zeta potential (ζ), were measured using DLS on a Zetasizer Nano-ZS instrument (Malvern Instruments, Ltd., UK). Measurements were taken using samples of MC NPs at a concentration of 1 mg/mL, and the average value was obtained from three measurements.

### Nanoparticle tracking analysis (NTA)

We conducted NTA using a ZetaView PMX 110 V3.0 instrument (Particle Metrix GmbH, Germany), and the data were analyzed with ZetaView NTA software. Samples were diluted 1:100,000 with PBS that had been filtered through a 20 nm syringe filter before analysis. We determined particle number, zeta potential, and size distribution at 11 positions with 2 acquisitions per position, under a sensitivity of 80 and a shutter value of 100.

### Transmission Electron Microscopy (TEM)

To monitor the morphology, we used TEM with an LEO EM910 TEM operating at 80 kV (Carl Zeiss SMT Inc., Peabody, MA). Diluted samples (100x) were placed on a copper/carbon grid and stained with 1% uranyl acetate before imaging.

### HPLC Analysis

To determine the concentration of Palb in MC NPs, we used reversed-phase high-pressure liquid chromatography (Agilent Technologies 1200 series) with a Nucleosil C18, 5 µm column (250 mm × 4.6 mm L × I.D.). The analysis procedures and settings were performed as previously described.^14^

### Spectroscopy

Fluorescence and UV-Vis absorption spectra were recorded using a FluoroLog spectrofluorometer (Jobin Yvon, Horiba) and Cary 4000 spectrophotometer (Varian), respectively. Absorbance measurements were performed at the maximum wavelengths (λ_max_) of 550 nm and 560 nm for RhB and NR dyes loaded in MC NPs. To determine their concentrations, fluorescence emission spectra were obtained at room temperature with excitation/emission (Ex/Em) wavelengths of 553/627 nm for RhB dye, and 562/655 nm for NR dye, and the spectra were recorded at 560–800 nm and 580–800 nm wavelengths, respectively. Fluorescence measurements were conducted using solutions with an absorbance of less than 0.2.

### Drug loading and release

NR and RhB dyes, as well as Palb MC formulations, were prepared at a concentration of 2 mg/mL, and their release rates from the NPs were measured under sink conditions in PBS at 37 °C. A 100 µL sample was placed in a 3.5 kDa floatable slide-A-lyzer Mini dialysis device, after washing the membrane with PBS to comply with perfect sink conditions. The samples were incubated in triplicate at 0, 0.5, 1, 2, 4, 8, and 24 h. At each time point, the solution was removed from the dialysis device, lyophilized, rehydrated, and diluted for analysis by either HPLC (for Palb) or fluorescence intensity (FI) measurements (for NR and RhB dyes), after centrifugation and removal of salts. We calculated the cargo loading efficiency (LE) and loading capacity (LC) using the following equations:

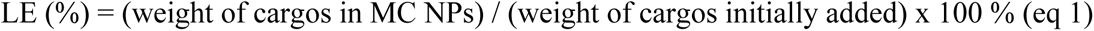

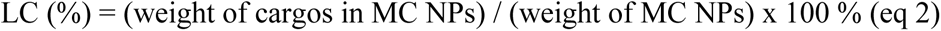

### Erythrocytes hemolysis assay

The hemolysis assay was conducted on a defibrinated sheep blood sample using a procedure that was previously described.^16^

### Size stability in FBS

A sample containing 100 µg/mL of MC NPs was incubated with FBS at 37 °C under gentle agitation for 48 h. The volume ratio of the NP solution to 100% FBS was 1:9. The size of the particles was measured using NTA at 0, 24, and 48 h time points. The data confirmed the high stability of the particles, as no significant changes in size were observed.

### Cell viability studies

Cell viability studies were conducted to evaluate the *in vitro* cytotoxicity of MC1 NPs, Palb-MC1 NPs, MC2 NPs, Palb-MC2 NPs, and free Palb on RAW 264.7 macrophage, U-87 GBM, and M14 melanoma cell lines. <u>The toxicity of MC NPs alone was also tested on HDFn and Mel 29.1 cell lines.</u> The cells were treated with various concentrations of all formulations, prepared by serial dilution in full medium, as indicated. Prior to treatment, cells were cultured and seeded in a 96-well plate at a density of 10^4^ cells/well and were allowed to attach for 24 h. Cell viability was assessed using the CCK-8 assay according to the manufacturer’s protocol at 24 and 72 h after treatment with all test articles. Data are presented as means ± SD of six replicate wells. Cytotoxicity measurements of MC1 NPs and MC2 NPs were also performed on adipose stromal cells by MTT assay according to previously described method.^17^

### Cell uptake studies

#### Fluorescence Imaging

U-87 MG cells were seeded in a 24-well plate at a density of 35,000 cells per well and allowed to attach overnight. The following day, cells were treated with either free NR or NR-loaded MC NPs for 4 or 24 h. After treatment, cells were washed, and NucBlue live cell stain was used to stain the nuclei. Fluorescent images were captured at 20x magnification using an EVOS FL microscope. The same treatment protocol was followed for flow cytometry. Cells were immediately trypsinized and removed from the plate for flow cytometry analysis, which was conducted using the BL3A laser on an Attune NxT to quantify NR dye uptake.

#### Confocal Laser Scanning Microscopy (CLSM) (Zeiss LSM 710) Imaging

Human M14 melanoma and human fibroblast HDFn cell lines were cultured in a 24 well-plate at a density of 5,000 cells per well and allowed to attach overnight. NR dye-loaded MC NPs at a concentration of 0.63 mg/ml were incubated in both cell lines for 8 hours. 1:1000 stain solution of CellMask Plasma Membrane stain was prepared in 1X PBS. Cells in suspension received 200 ul of stain solution, were covered in foil, and left in the dark for 10 minutes. Cells were then washed with 1X PBS and spun at 400G for 5 minutes. Stain cells were suspended in Fix & Perm Buffer with Media. PRMI was used for both M14 and HDFn cells.

### Flow cytometry for quantification of live/dead cells

To quantify live/dead cells, we conducted flow cytometry evaluations using GFP expression on tumor cells and mouse macrophages F4/80 conjugated with phycoerythrin (PE). Samples were acquired using either BD FACSCanto II or BD LSRFortessa and analyzed with BD FACSDiva software (BD Biosciences). We evaluated the quantification of cells treated with free Palb, blank MC NPs, and Palb-loaded MC NPs at two concentrations and compared them to untreated cells. A minimum of 50,000 events were acquired for each sample, and FlowJo version 8 was used for data analysis.

## 3. Results and discussion

The two compounds utilized in our study, MC1 and MC2, are composed of two long alkyl chains attached to the MC core. MC1 contains a nitro group, while MC2 bears an amino group (**Figure 1a**). Our objective was to formulate NPs stabilized by CrEL using MC1 and MC2. First, we demonstrated the ability of the resulting MC-based NPs to incorporate hydrophilic and hydrophobic fluorescent dyes as model drugs. Second, we optimized the MC-based formulations to improve the delivery of the CDK4/6 inhibitor, Palbociclib (Palb), for melanoma treatment. To prepare MC NP formulations, we used the solvent diffusion method (**Figure 1b**), where all ingredients (drug, MCs, and CrEL) were dissolved in a water-miscible solvent and added dropwise to an aqueous phase. The organic solvent and unincorporated solutes were removed by centrifugal filtration (10 kDa). We varied the water-miscible solvent (DMSO or THF) and the surfactant to MCs mass ratio to explore multiple conditions (**Figure 1c**). Without CrEL, the MCs tend to spontaneously self-assemble into NPs, but they can only remain stable at relatively low concentrations, up to ∼600 µM (∼0.6 mg/mL). Our results indicated that the addition of CrEL stabilized the MC NPs in dispersion up to ∼10 mM (∼10 mg/mL) of MCs. The optimal formulation was obtained at 36 and 64 wt. % of MCs and CrEL, respectively. For all subsequent experiments, we used this condition to describe the surfactant-stabilized NPs. We then loaded NR dye (hydrophobic) and RhB dye (hydrophilic) into these optimal formulations and obtained colored dispersions, as shown in **Figure 1d**.

**Figure 1.**
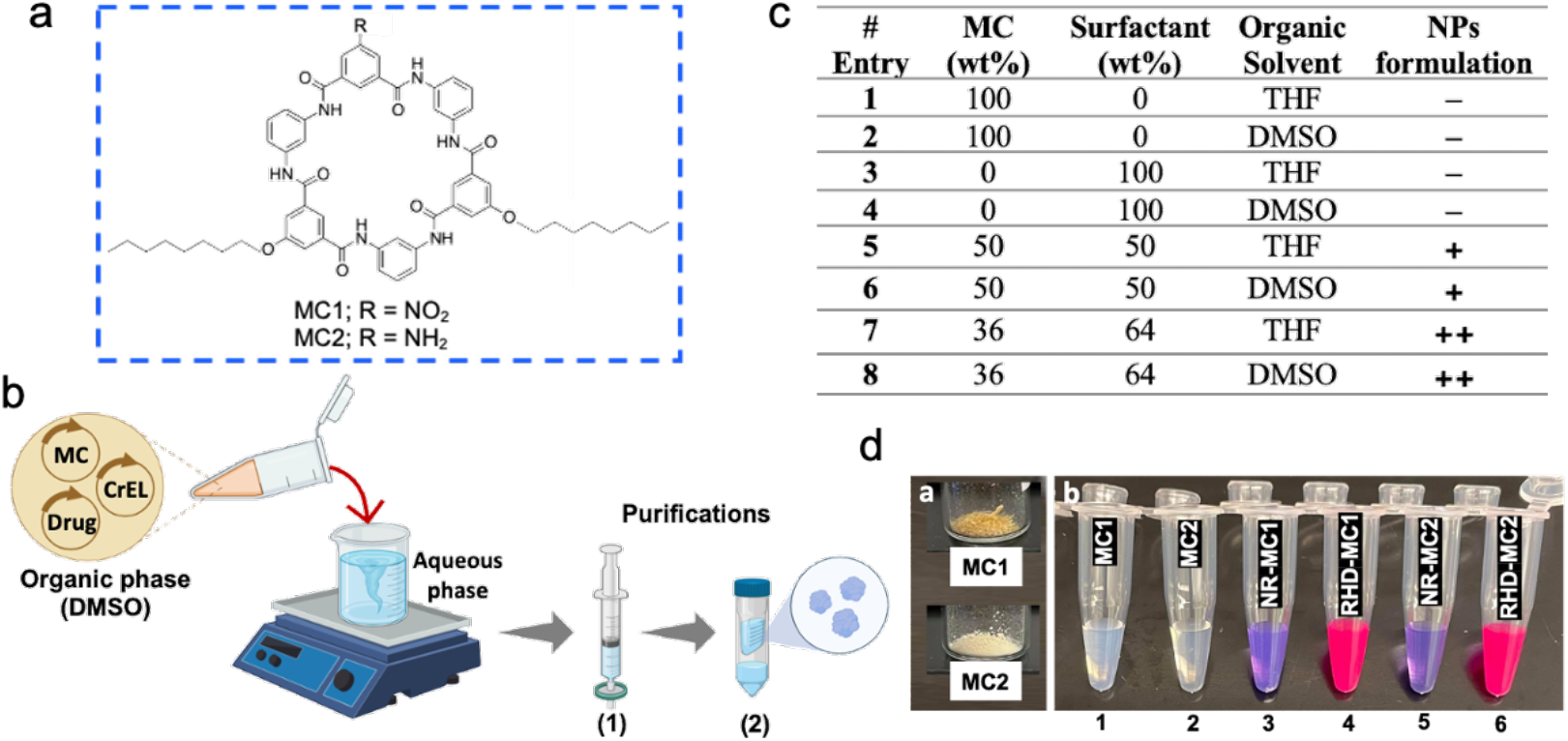
**(a)** The chemical structure of MC compounds, **(b)** the formulation and purification processes of MC NPs by solvent diffusion method, and **(c)** the optimization of nanoformulations using different MC/surfactant weight ratios and organic solvents (THF and DMSO). Formulations were rated as (−) bad, (+) good, and (++) excellent based on particle size and PDI. Additionally, **(d)** digital photos of MC1 and MC2 materials (left) and MC1 and MC2 NPs aqueous dispersions (1-6) stabilized by PEGylated surfactant at the weight ratio of 36:64, respectively, are shown. Tubes 1, 3, and 4 represent MC1 NPs, NR dye-loaded MC1 NPs, and RhB dye-loaded MC1 NPs, respectively, while tubes 2, 5, and 6 represent MC2 NPs, NR dye-loaded MC2 NPs, and RhB dye-loaded MC2 NPs, respectively.

TEM analysis of non-formulated MC1 and MC2 materials, which were deposited from THF dispersion, revealed heterogeneous populations of spherical aggregates with particle sizes ranging from approximately 120 to 350 nm (**Figure 2a,e**). Similar aggregates have been reported previously.^8^ In contrast, the surfactant-stabilized MC NPs dispersed in water appeared to be smaller and more uniform, although they exhibited a tendency to stack in a toy pyramid fashion (**Figure 2b,f**). The DLS analysis of these surfactant-stabilized NPs showed relatively narrow size distributions (PDI < 0.2) with effective particle diameters of 106 nm and 122 nm for MC1 and MC2 NPs, respectively (**Figure 2c,g**). Interestingly, the size distributions and diameters of these particles were very similar to particles prepared using both DMSO and THF in the formulation process. The dye-loaded formulations of NR-MC1 and RhB-MC2 NPs were also imaged by TEM (**Figure 2d,h**). Notably, their size and morphology were similar to those of the blank surfactant-stabilized NPs.

**Figure 2.**
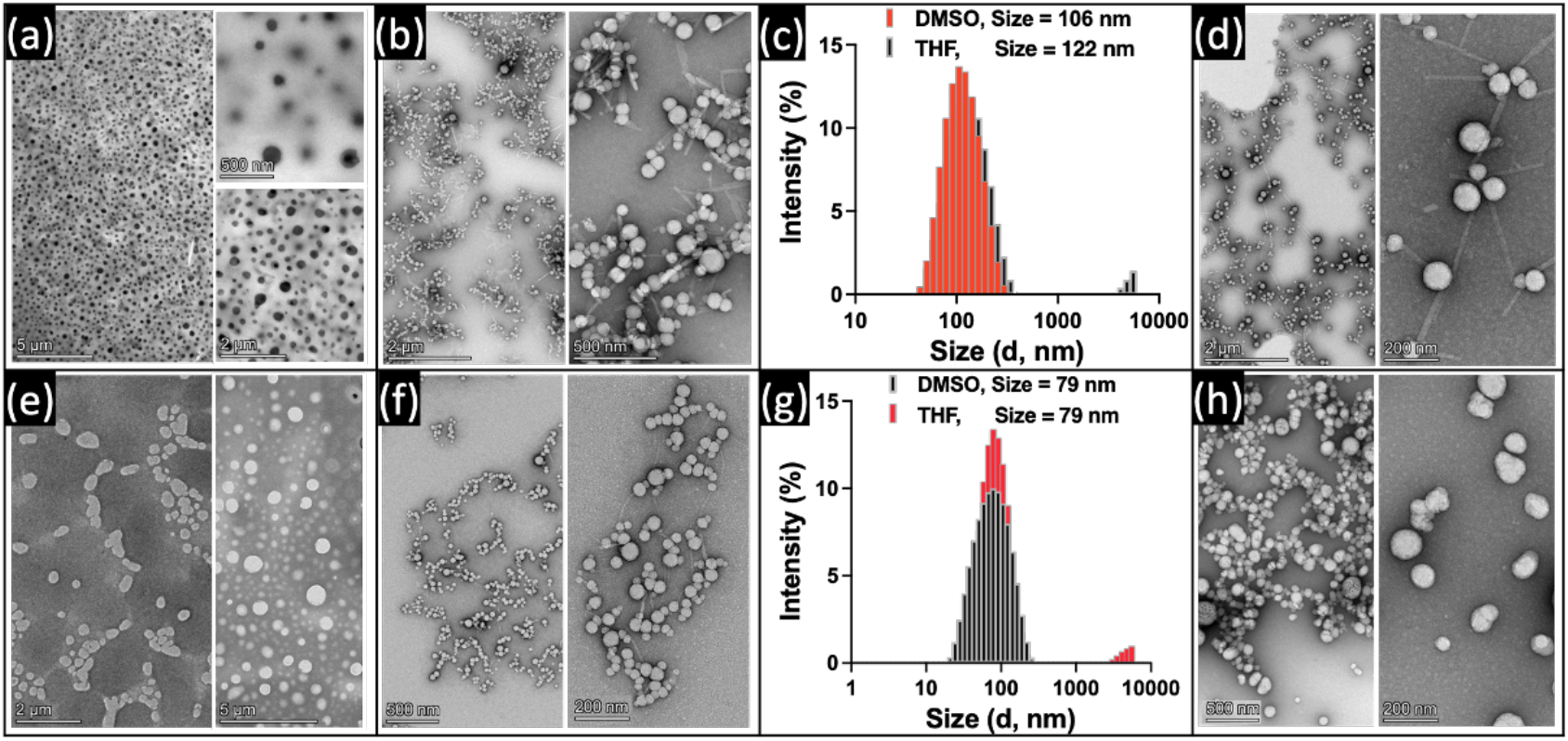
The characterization of (a-d) MC1- and (e-h) MC2-based materials. TEM images of **(a, e)** pure MC1 and MC2 dried on a grid from THF, **(b, f)** unloaded MC1 and MC2-based NPs (both prepared in DMSO), and **(d, h)** NR-MC1 NPs and RhB-MC2 NPs (both prepared in DMSO) are presented at two different magnifications. Additionally, the intensity particle size distribution of empty MC1 and MC2 NPs (prepared in either THF or DMSO) is shown using DLS in (c, g).

We confirmed the incorporation of the dyes into the MC NPs by analyzing the fluorescence spectra of free and dye-loaded MC NPs (**Figure 3**). We observed the appearance of peaks corresponding to the dye fluorescence in both dyes loaded into MC1 or MC2 NPs, which were not present in the case of empty MC NPs.

**Figure 3.**
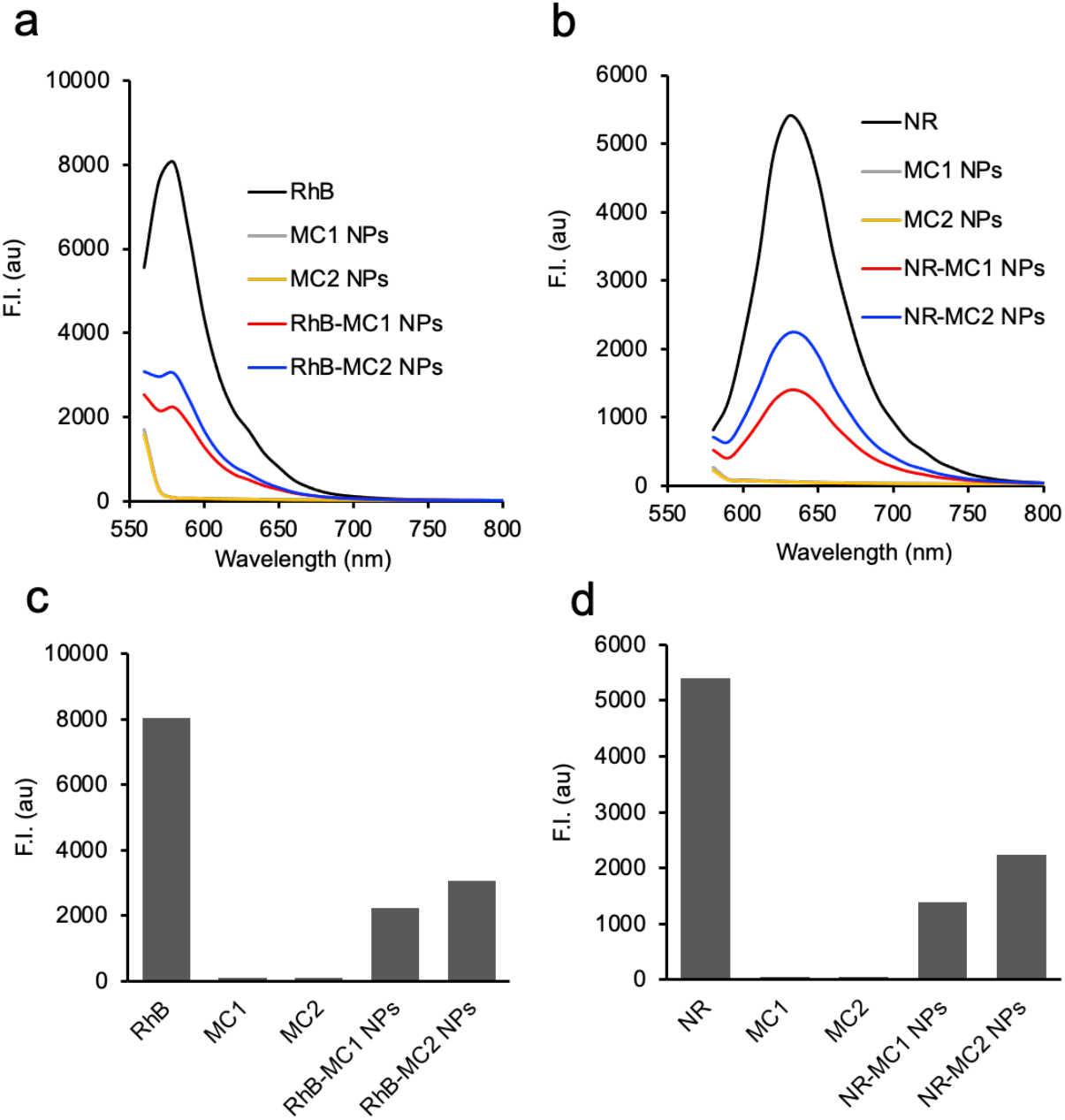
Fluorescence emission (λex 562 nm) spectra of MC NP loaded with (a) RhB and NR (b) dyes. All samples were prepared at MC (1 mg) : PEG (1.8 mg) : dye (1 mg) in a final volume of 1 mL of bi-distilled water. The excess of unloaded dyes was separated by ultracentrifugation. For the spectroscopic measurements. The spectra were recorded at the following final concentrations of the dyes (see Supplementary Table S1).

Next, we calculated the loading efficiency (LE) and loading capacity (LC) of both dyes in the MC1 and MC2 NP formulations by means of fluorescence intensity measurements. Calibration curves were made for free dyes to determine the endpoint fluorescence in terms of dye concentrations at the same Ex/Em for each dye as mentioned above (**Figure 4a,b**). The results showed that the loading efficiency (LE) for RhB-MC1 was 17%, whereas it was slightly higher at 21% for RhB-MC2. Similarly, NR-MC2 exhibited a higher LE (23%) than NR-MC1 (12%). We also calculated the loading capacity (LC), which was found to be 6-8.7% for RhB in MC1 and MC2, respectively, and 4-8% for NR in MC1 and MC2, respectively (**Figure 4c**). Overall, these findings suggest that MC2 demonstrated higher entrapment efficiency as compared to MC1, likely due to the potential for favorable hydrogen bonding of the dyes with the amino groups of MC2.

**Figure 4.**
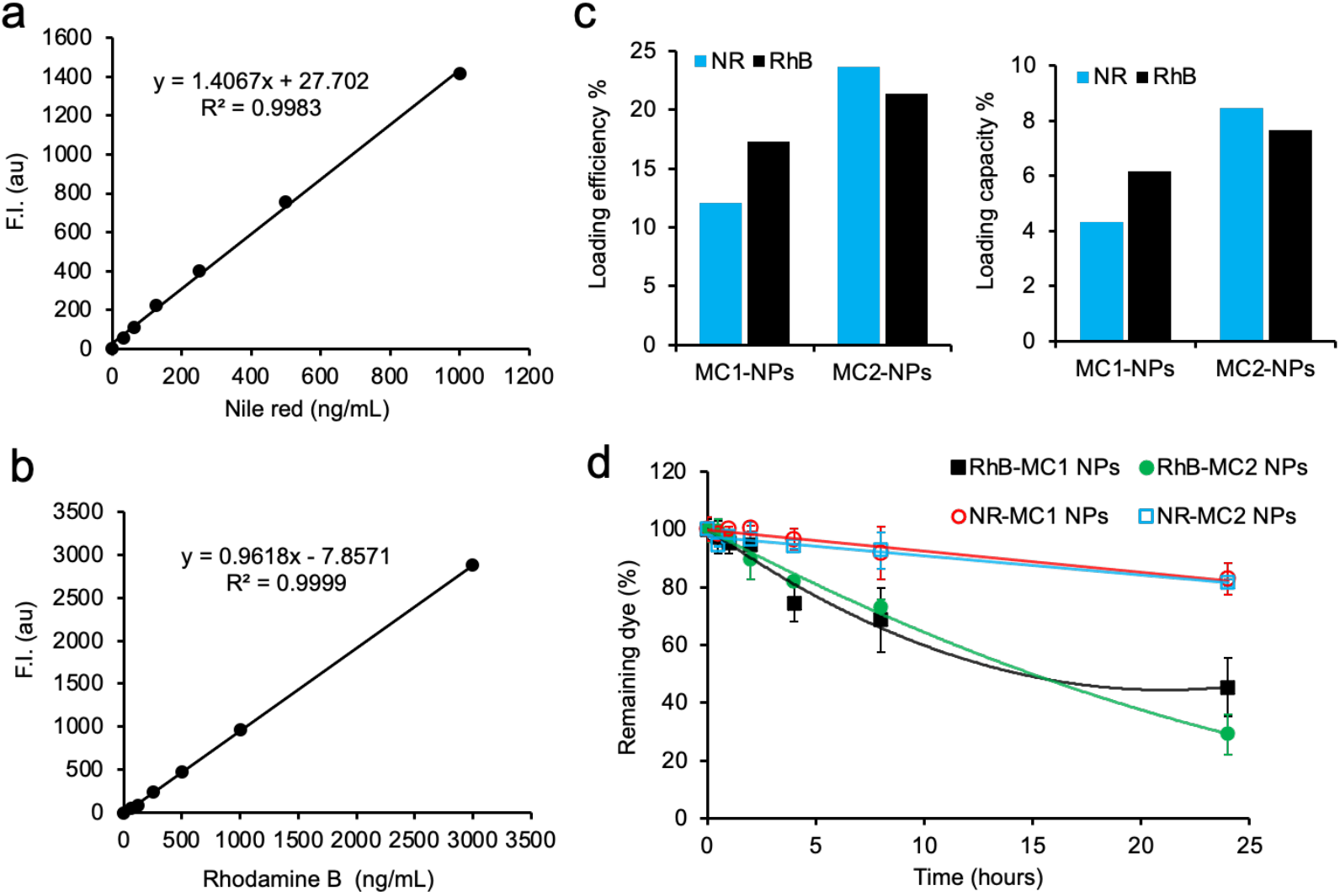
(a,b) Fluorescence calibration curves of free NR and RhB, respectively (RhB Ex/Em 553/627 nm; NR Ex/Em 562/655 nm). (c) Dye loading capacities (LC) and loading efficiencies (LE) in MC NPs. (d) Dye release kinetics from MC NPs.

A highly desirable feature for drug delivery nanocarriers is slow and controlled release kinetics. In our study, we observed that the release kinetics strongly depended on the structure of the dye. However, there was a similarity between the release profiles of each dye in both types of MC NPs, as shown in **Figure 4d**. We found that both MC NPs released between 55% to 70% of RhB over 24 h. In contrast, the release of NR from both MC NPs was much slower, with only about 20% of the dye released over the same period. This difference in release kinetics is likely due to the interaction of the dyes with the MC NPs. The hydrophobic NR molecule was probably bound in the cavities of MC NPs, while the more hydrophilic RhB was more likely to interact with the outer shell of MC NPs. Overall, both MC NPs exhibited slow and controlled release profiles compared to polymeric and liposomal NP/drug conjugates.

We monitored the particle size of all blank and loaded MC NPs immediately after formulation and after one month using NTA Zetaview to assess their stability over time. The data showed that the materials had a typical narrow size distribution both before and after one month of preparation, with negligible change in size (**Figure 5a**, and Supplementary **Figure S1**). It should be noted that the net negative surface charge increased over time (**Figure 5b**), and there was no noticeable change in particle concentration as determined by NTA (**Figure 5c**). We also tested the stability of the materials in the presence of serum by recording the size of all MC NPs at 0, 24, and 48 h time points (**Figure 5d**). This experiment demonstrated that the materials maintained a very stable particle size without any aggregation upon incubation in FBS.

**Figure 5.**
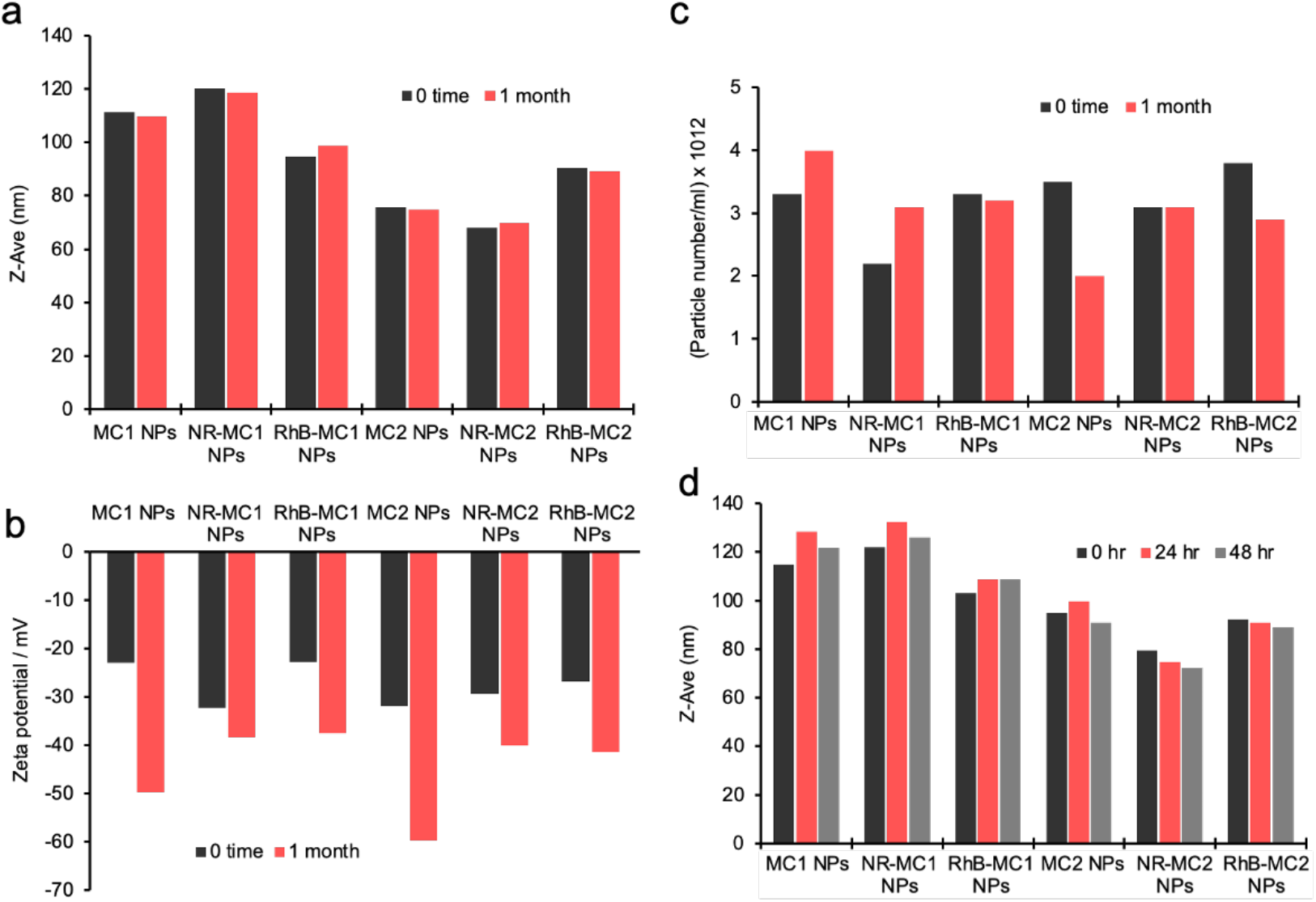
The results of NTA of the blank and dye-loaded MC NPs. **(a)** Mean particle size of all nanoparticles immediately after preparation and after one month, indicating no variation in size. **(b)** Zeta potential, **(c)** particles’ concentration, and **(d)** stability of MC NPs in FBS over 48 h. The stock NPs solution was diluted 10-fold by FBS.

Next, we prepared drug-loaded MC NPs by encapsulating a hydrophobic drug, Palb (free base), and evaluated their size, zeta potential, morphology, LC, LE, and hemolysis profiles. We prepared the nanoformulations using DMSO as the solvent with an MC : CrEL : Palb weight ratio of 2 : 3.6 : 0.5, following the same protocol as above. The resulting Palb-loaded MC NPs exhibited diameters of 90 nm and 64 nm for Palb-MC1 and Palb-MC2 NPs, respectively, with a narrow size distribution as determined by NTA (**Figure 6a,b**). The complexes had highly negative zeta potentials of –28 and –39 mV for Palb-MC1 and Palb-MC2 NPs, respectively (**Figure 6b,e**). TEM images showed uniform, spherical particles within the same size range observed in the NTA data (**Figure 6c,f**).

**Figure 6.**
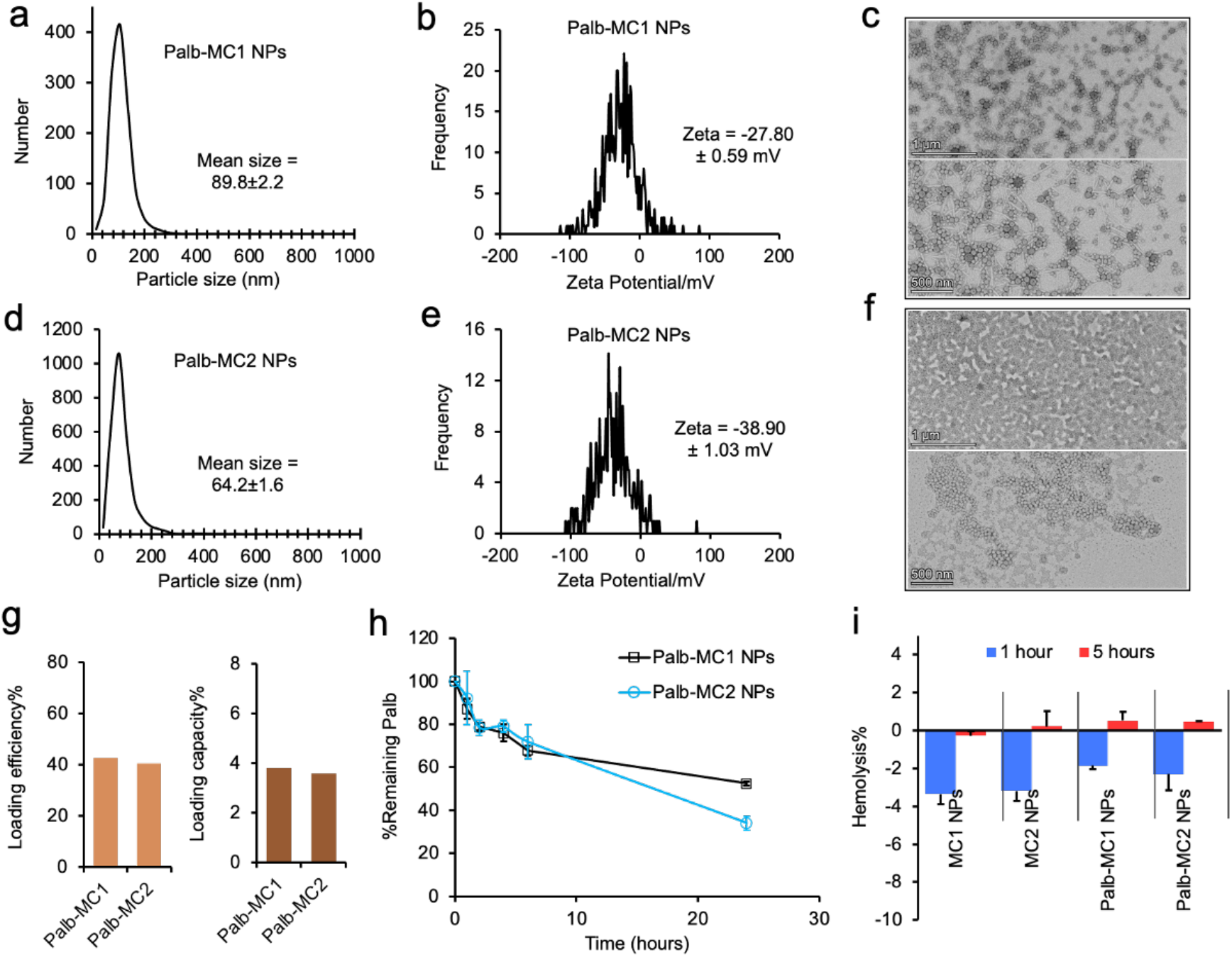
Characterization of Palb-loaded MC NPs. **(a, d)** Particle size distribution, **(b, e)** zeta potential (measured by NTA), and **(c, f)** TEM images of Palb-MC1 and Palb-MC2 NPs, respectively. **(g)** LC and LE percentages of Palb in MC1 and MC2 NPs. **(h)** Palb release profile from MC1 and MC2 NPs. **(i)** Hemolysis assay results for unloaded and Palb-MC NPs.

The LC and LE profiles of the MC-Palb NPs were analyzed by HPLC, revealing approximately 40% LE for both MC1 and MC2 NPs, with nearly identical LCs of 4% (**Figure 6g**). The release kinetics demonstrated a slow and controlled release of Palb from both MC NPs, with approximately 28% of the drug being released from MC NPs after 6 h (**Figure 6h**). The total release of the drug from MC1 and MC2 NPs after 24 h was 47% and 66%, respectively.

The *in vitro* hemolytic properties of both MCs and Palb-MC NPs were evaluated using red blood cells (erythrocytes) collected from sheep. Spectrophotometric measurements were taken after incubating the MCs with the blood cells for 1 and 5 h, as described in detail in the methods section. The results showed that the hemolysis induced by MCs was negligible and did not cause any cell lysis or death during the assay period, indicating that the materials are cytocompatible (**Figure 6i**). To assess the biocompatibility of blank MC NPs, we evaluated their cytotoxicity against adipose stem cells (shown in Supplementary **Figure S2**). After 24 h of treatment, both MC NPs demonstrated 100% cell viability at a concentration of 500 µg/mL. At a higher concentration of 3000 µg/mL, the viability was reduced to nearly 60%. These findings suggest that the MC1 and MC2 NPs exhibit excellent biocompatibility.

To confirm the biocompatibility of MC NPs, we evaluated the viability of RAW264.7 macrophage cells treated with various concentrations of blank MCs, Palb-MC NPs, and free Palb using a CCK-8 assay for 24 and 72 <u>hours</u> (**Figure 7**). Both MCs showed excellent biocompatibility at higher concentrations (up to 1.26 mg/mL), as suggested by 100% viability <u>after</u> 24 <u>hours exposure</u> (refer to Supplementary **Figure S3** for details). However, increasing the exposure of the cells to our materials to 72 hours resulted in increased cytotoxicity. Nonetheless, cells were 100% viable up to 0.63 mg/mL MCs. At 24 hours of exposure, free Palb was not toxic to cells up to 6.25 µM; however, at 12.5 µM, it led to a decrease in <u>cell</u> viability to 20%. After <u>72 hours, cells treated with up to 3.12 µM</u> free Palb displayed 100% viability, while at 6.25 µM and above, there was a clear cytotoxic effect of the drug. After 72 hours of treatment, Palb loaded in MC1 and MC2 NPs exhibited slightly higher toxicity than the free drug. This is evident when comparing these groups with the free Palb treatments ranging from 1.56 µM to 6.25 µM. This result may be attributed to the improved intracellular delivery of the drug facilitated by the MC NPs. (Supplementary **Table S2** represents IC50 values for MC NPs and Palbociclib drug for both exposure times in RAW264.7 cells).

**Figure 7.**
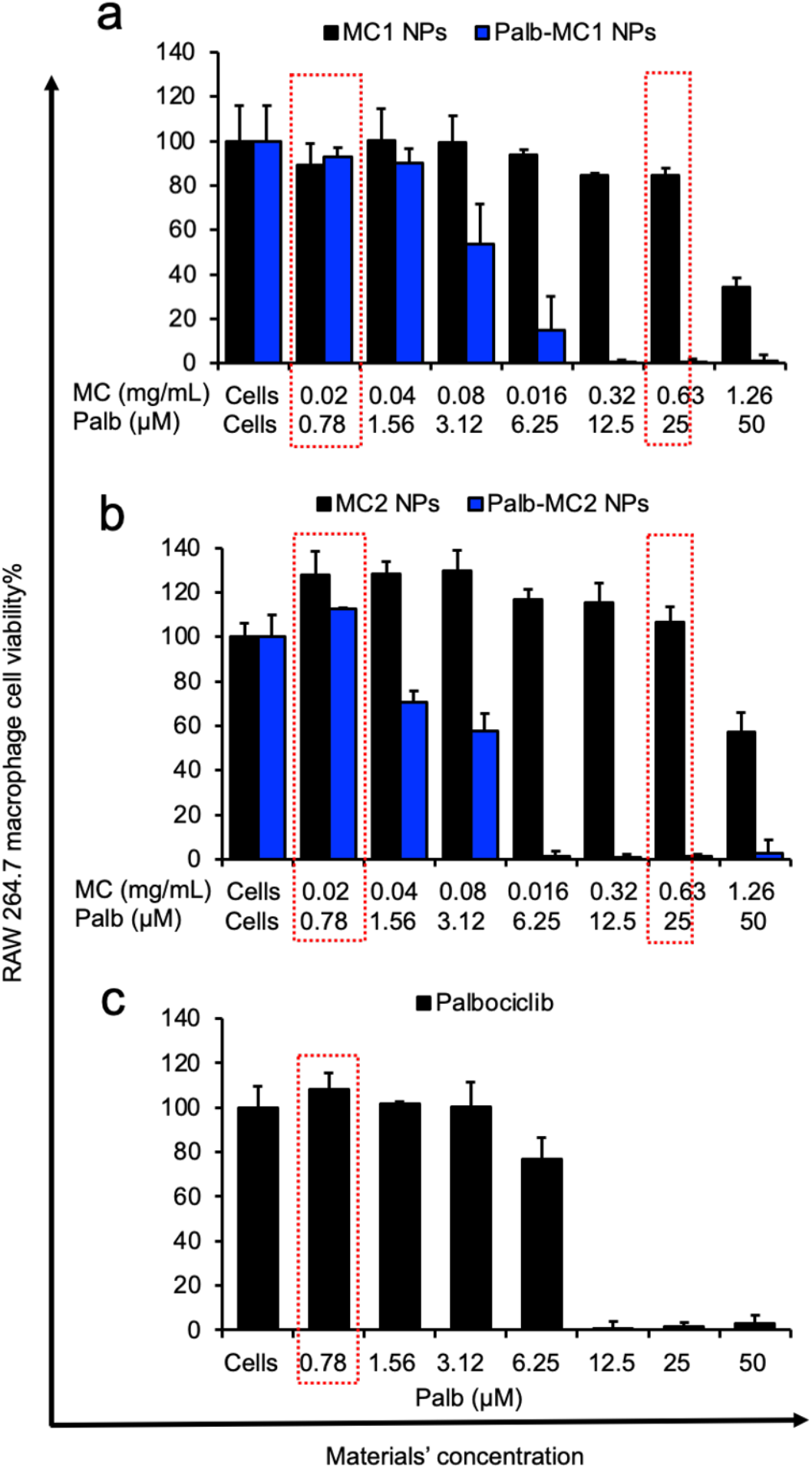
Cell viability studies (measured by CCK-8 assay) <u>using</u> RAW264.7 macrophage cells treated with various concentrations of blank MC NPs, Palb-MC NPs, and free Palb drug. The cells were exposed to the test nanoparticles for 72 h, and the results of the 24 h exposure are presented in Supplementary **Figure S3**. The highlighted treatments in red rectangles were selected for flow cytometry analysis to quantify cell death caused by different treatments. Viability of cells treated with **(a)** blank MC1 NPs Palb-MC1 NPs. **(b)** blank MC2 NPs Palb-MC2 NPs, and **(c)** free Palb drug.

<u>Next, we examined the effects of MC NPs with and without the drug in</u> RAW264.7, U-87 GBM-GFP, and M14-GFP melanoma cell lines <u>under a fluorescent microscope</u> to monitor <u>cell</u> morphology and proliferation (**Figure 8**). In this experiment, we used two concentrations of MC NPs, 0.02 mg/mL and 0.63 mg/mL that were shown to be safe to our macrophage cells. With all treatments, we observed no differences in cell morphology and cell density between the treated and untreated RAW264.7 and GBM cells. However, in the case of M14 human cells, we noticed clear evidence of cell death upon treatment with <u>both</u> blank <u>MCs</u> and Palb-MC NPs at a concentration of 0.02 mg/mL of MC NPs. Additionally, significant cell death was observed in M14 cells treated with MC only at higher concentrations of 0.63 mg/mL. These results indicate that both unloaded, and Palb-loaded MC NPs <u>can</u> kill melanoma cells, as compared to cells treated with free Palb and untreated controls.

**Figure 8.**
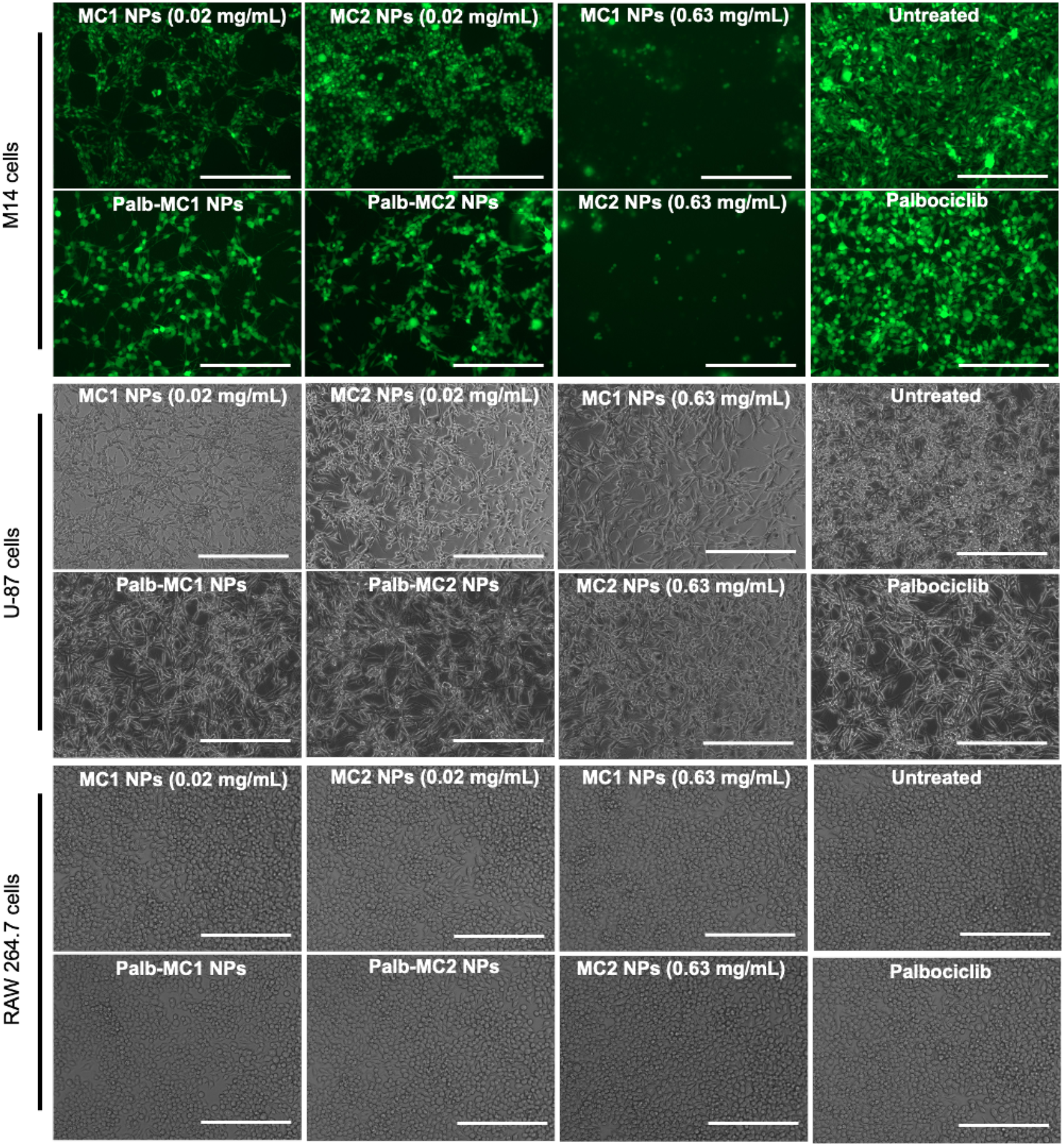
Fluorescence microscopy images of M14-GFP melanoma, U-87 GBM-GFP, and RAW264.7 macrophage cells after 72 h of treatment, demonstrating the influence of the type and concentrations of MC NPs on cell death. The concentration of MC1 and MC2 in either drug-free or Palb-containing treatment solutions was either 0.02 mg/ml or 0.63 mg/ml as indicated. The Palb drug was used at a concentration of 0.78 µM as free or MC-formulated solution. The colored images show M14-GFP melanoma cells treated with different concentrations of MC NPs, as indicated in each image, while the black-and-white images depict RAW264.7 macrophages and U-87 GBM-GFP cells treated with the same concentrations. A scale bar of 200 µm is present in all panels.

To measure the differential therapeutic effect of MC NPs on melanoma cells versus macrophage and GBM cells, we used flow cytometry in the same experiment. The flow cytometry data (**Figure 9**) was consistent with the fluorescence microscopy images above, indicating that none of the treatments caused obvious cell death in U-87 GBM-GFP and RAW264.7 macrophage cells. However, treatment of M14-GFP melanoma cells with MC NPs (0.63 mg/mL), Palb-MC NPs <u>(0.02 mg/mL)</u>, and free Palb (0.78 µM) resulted in severe toxicity, with 70-80% of all cells undergoing cell death.

**Figure 9.**
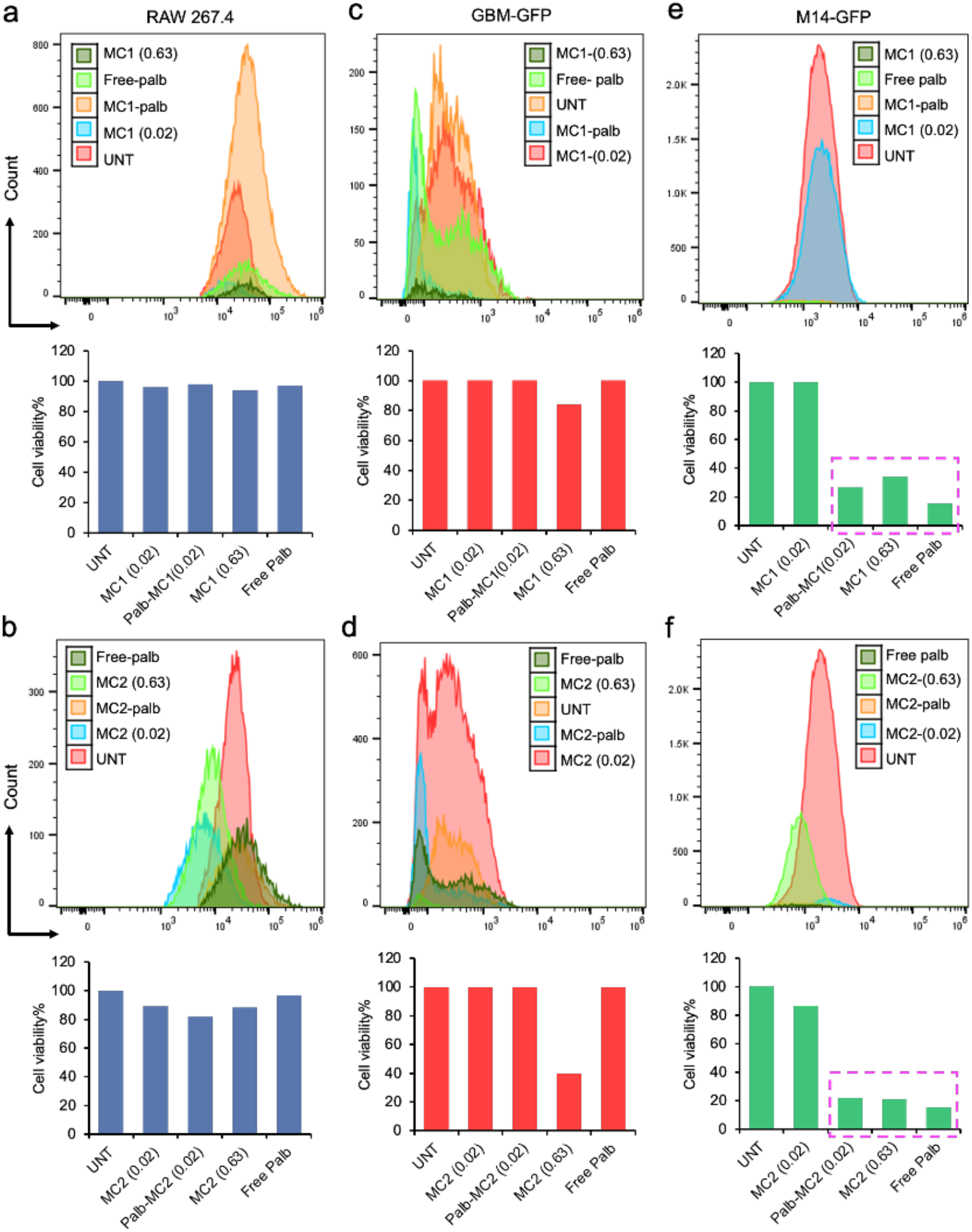
Representative flow cytometry profile and comparative cytotoxic effect after 72 h of treatment, demonstrating the selective killing of melanoma cells by MC1 and MC2 NPs. Panels **(a)** and **(b)** show the treatment of macrophages RAW264.7 cells with MC1 and MC2 NPs, while panels **(c)** and **(d)** show the treatment of U-87 GBM-GFP glioblastoma cells. Panels **(e)** and **(f)** depict the treatment of M14-GFP melanoma cells. The drug doses were the same as described in Figure 8.

This unusual result suggests a “specificity” of the cytotoxic effect in melanoma, indicating that this effect is attributed not to the anticancer drug but to MC NPs. To validate this assumption, we examined the effects of MCs at different concentrations on several cell lines, including Human Dermal Fibroblasts (HDFn), U-87 GBM-GFP, and two melanoma cell lines, M14-GFP and Mel 29.1 (**Figure 10**). Like what was observed in RAW264.7 macrophages in the fibroblast culture, MCs did not show any toxicity up to 0.63 mg/ml after 24 hours and 72 hours of exposure (**Figure 7**, supplemental **Figure S5**). In contrast, in both melanoma cell lines, MCs exhibited significant cytotoxicity at much lower concentrations (**Figure 10**). The greatest cytotoxic effect was observed in Mel 29.1 cells, with estimated IC50 values for MC1 and MC2 being 0.09 mg/mL and 0.16 mg/mL, respectively (see supplementary **Table S3**). A similar pattern was observed in M14-GFP cells, with MC1 being slightly more toxic (IC50 0.22 mg/mL) than MC2 (IC50 0.43 mg/mL). The MCs also exhibited cytotoxicity in the GBM cell line, particularly MC1, which displayed an IC50 of 0.26 mg/mL, whereas MC2 had an IC50 of 1.01 mg/mL.

**Figure 10.**
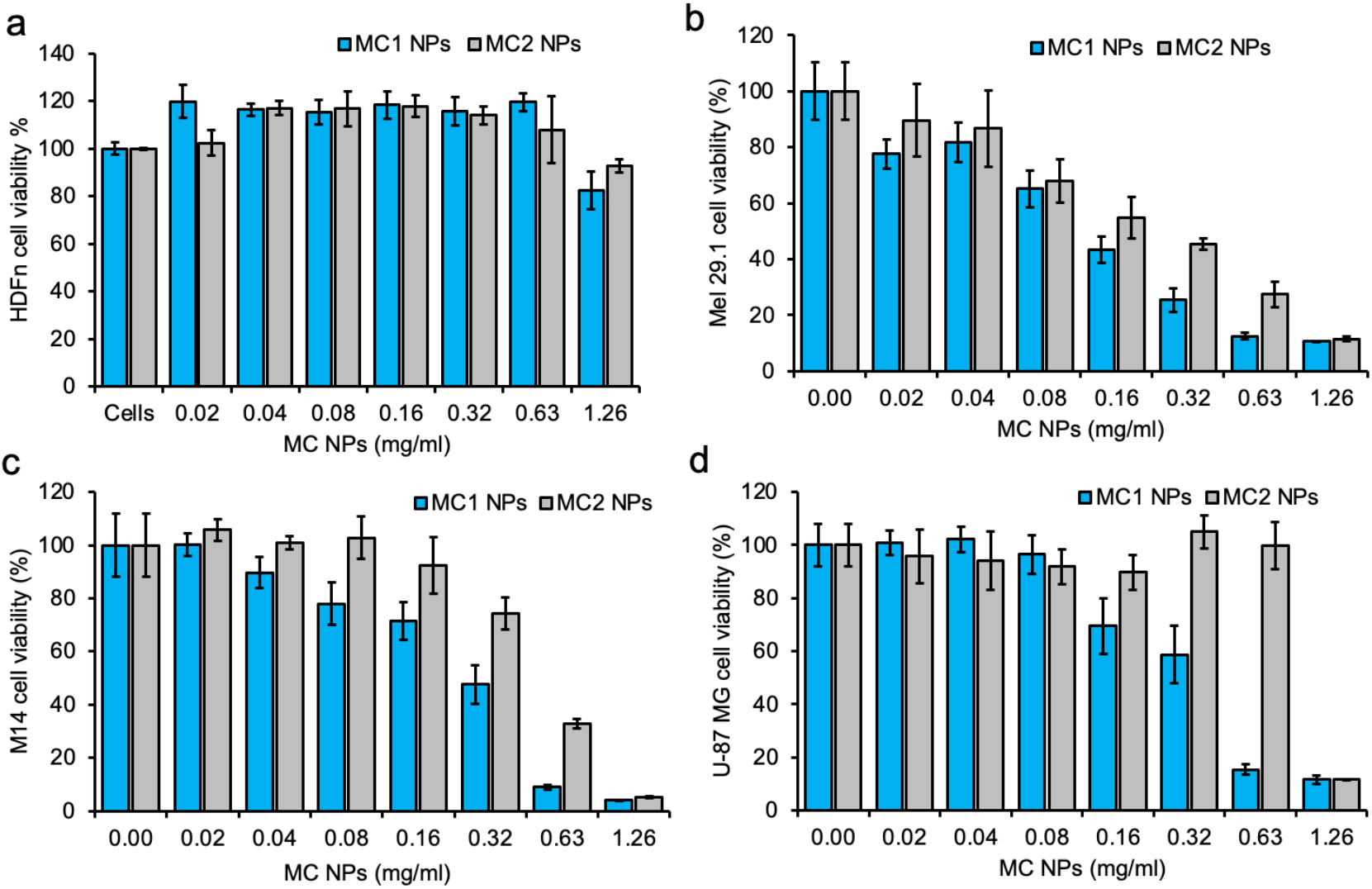
Cell viability of (a) HDFn), (b) Mel 29.1, (c) M14, and (d) U-87 MG cells treated with MC NPs for 72 hours of incubation. The experiment was done in triplicates.

To investigate the cellular uptake of MC NPs and their ability to internalize into the cytoplasm of U-87 GBM cells, we used NR dye as a tracer. We encapsulated NR dye in both MC NPs and compared the internalization of NR-MC NPs to positive (free NR) and negative controls (cells with unloaded MC NPs) after 4 and 24 h of incubation using fluorescence microscopy (see Supplementary **Figure S4**). The images showed a strong red fluorescence signal both after 4 and 24 h of incubation, indicating that both MC NP formulations were effectively internalized. Flow cytometry data confirmed that U-87 GBM cells treated with both NR-MC1 and NR-MC2 exhibited high levels of cellular uptake (approximately 70%) compared to free NR. These findings demonstrate the ability of MC NPs to efficiently enter cancer cells and deliver their cargo.

In a separate study, we tracked the internalization of NR-loaded MC NPs into human fibroblast HDFn and human melanoma M14 cells to investigate the reason for the different toxicity profiles. Confocal microscopy (**Figure 11**) showed that both MC NPs had similar cellular uptake and localized in the cytoplasm of both cells. Future studies will need to investigate the mechanistic reason for the apparent increased toxicity of the plain MCs to melanoma cells and whether this toxicity relates to specific interactions of our new materials with melanin. Literature suggests that melanin-binding drugs are more toxic to cells with high melanin content.^18–21^

**Figure 11.**
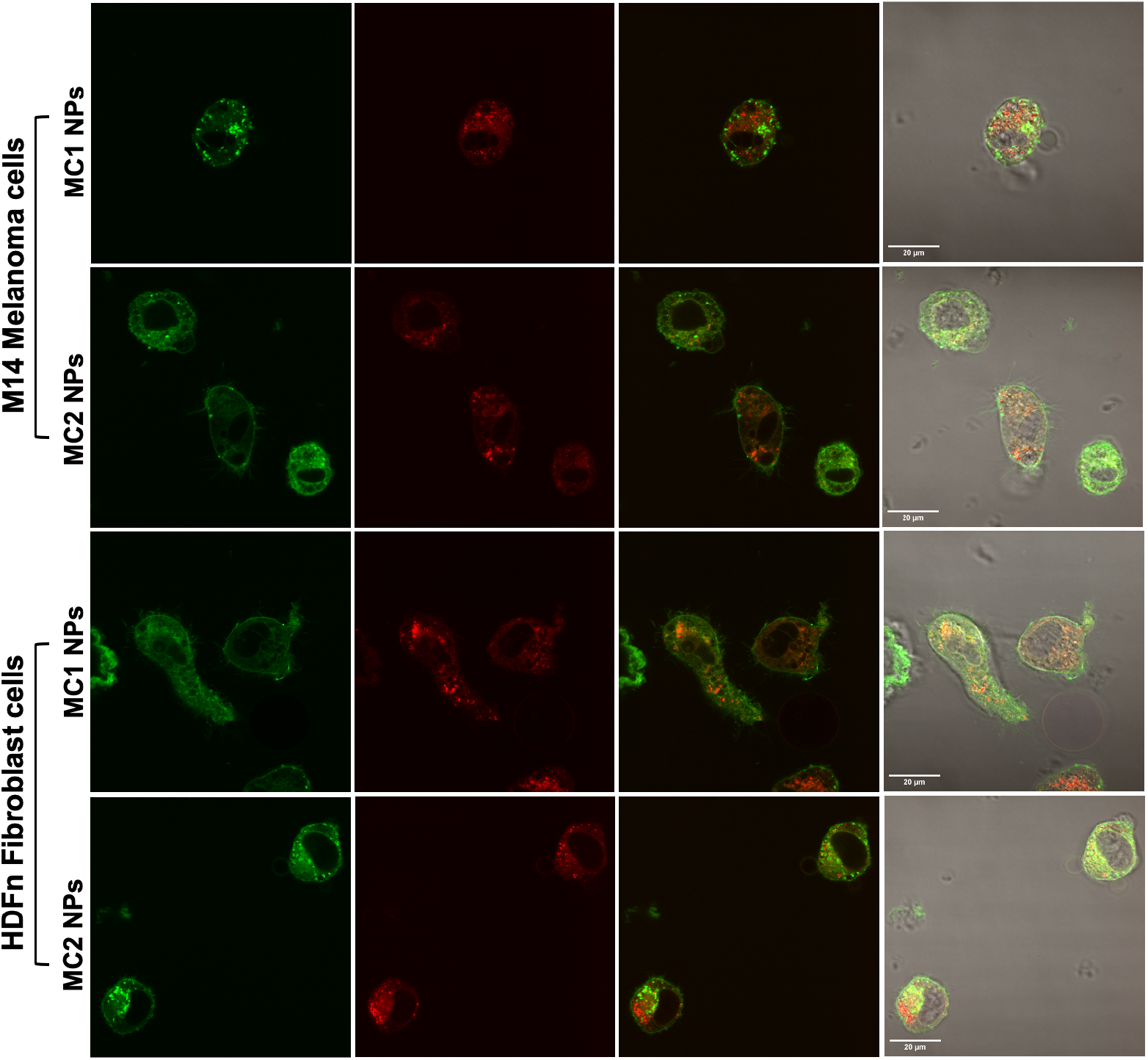
Confocal microscopy imaging of NR-MC1 and NR-MC2 NPs internalization in both M14 melanoma and HDFn fibroblast cell lines. The scale bar is 20 µm. Images in each row (left to right): CellMask (green) for staining cell membrane, NR-loaded MC NPs (red), overlapping (green-red), and bright field image (green-red).

## 4. Conclusion

In this study, we synthesized two new amphiphilic macrocycles and explored their potential as a drug delivery platform. We developed nonionic PEGylated surfactant-stabilized MC NPs that can incorporate both hydrophilic and hydrophobic molecules. By loading the CDK4/6 inhibitor Palb into these MC NPs, we improved drug delivery characteristics. We conducted physicochemical characterizations on both blank and Palb-MC NPs, which demonstrated high particle stability in serum, excellent biocompatibility, and a sustained, controlled drug release profile. <u>Moreover, our cytotoxicity and flow cytometry data suggest that both unloaded MCs and Palb-MC NPs exhibit a more pronounced cytotoxic effect in two melanoma cell lines without inducing cell death in macrophage and fibroblast cells at the same concentrations *in vitro*.</u> These promising results encourage testing the therapeutic activities of these materials in melanoma-bearing mice. Future experiments will focus on a comparative study to evaluate the long-term stability, toxicity, PK and biodistribution of the nanoformulated drugs. We will also investigate the potential of these materials to improve the water solubility and overall delivery of a library of hydrophobic chemotherapies.

## Conflict of interest

There are no conflicts to declare.

## Acknowledgments

MFA received support from the Carolina Cancer Nanotechnology Training Program, which is funded by NCI’s grant T32CA196589. EAO was supported by a grant from the National Institutes of Health K99NS128716. The research conducted in the Kabanov lab was partially supported by the National Cancer Institute (NCI) grant CA264488. The Whitehead laboratory is partially supported by NIH NIGMS grant 1P20GM146584-01. We thank Dr. Gianpietro Dotti (Lineberger Comprehensive Cancer Center, Chapel Hill, NC, USA) for providing GBM-GFP and M14-GFP cells for this study. We utilized ChatGPT 3.5 <u>and Microsoft Bing Chat Enterprise</u> as editorial tools to enhance the clarity and style of our fully drafted scientific manuscript, while also ensuring that the scientific content and integrity were preserved. While <u>AI chatbots</u> proved to be useful resources, we carefully reviewed and evaluated all suggested edits to ensure they aligned with our intended meaning and scientific accuracy. Our goal was to present our research findings in the most clear and accurate manner possible, and we believe that AI chatbots assisted us in achieving that goal.

## Notes

### Competing Interest Statement

The authors have declared no competing interest.

