## Supplementary information for "Enhancing Drug Delivery with Supramolecular Amphiphilic Macrocycle Nanoparticles: Selective Targeting of CDK4/6 Inhibitor Palbociclib to Melanoma"

**Affiliations:**

Table S1: Normalized fluorescence of RhB and NR dyes. (RhB Ex/Em 553/627 nm; NR Ex/Em 562/655 nm).

| **Samples** | **FI** | **Concentration (µM)** |
|---|---|---|
| RhB | 8047.585 | 17.5 |
| RhB -MC1 | 2234.446 | 4.9 |
| RhB -MC2 | 3043.415 | 6.6 |
| NR | 5396.946 | 12 |
| NR-MC1 | 1391.912 | 4.3 |
| NR-MC2 | 2231.056 | 6.9 |

Figure S1. Particle size distribution (measured by NTA Zetaview analysis) of all formulations immediately after preparation and after one month.

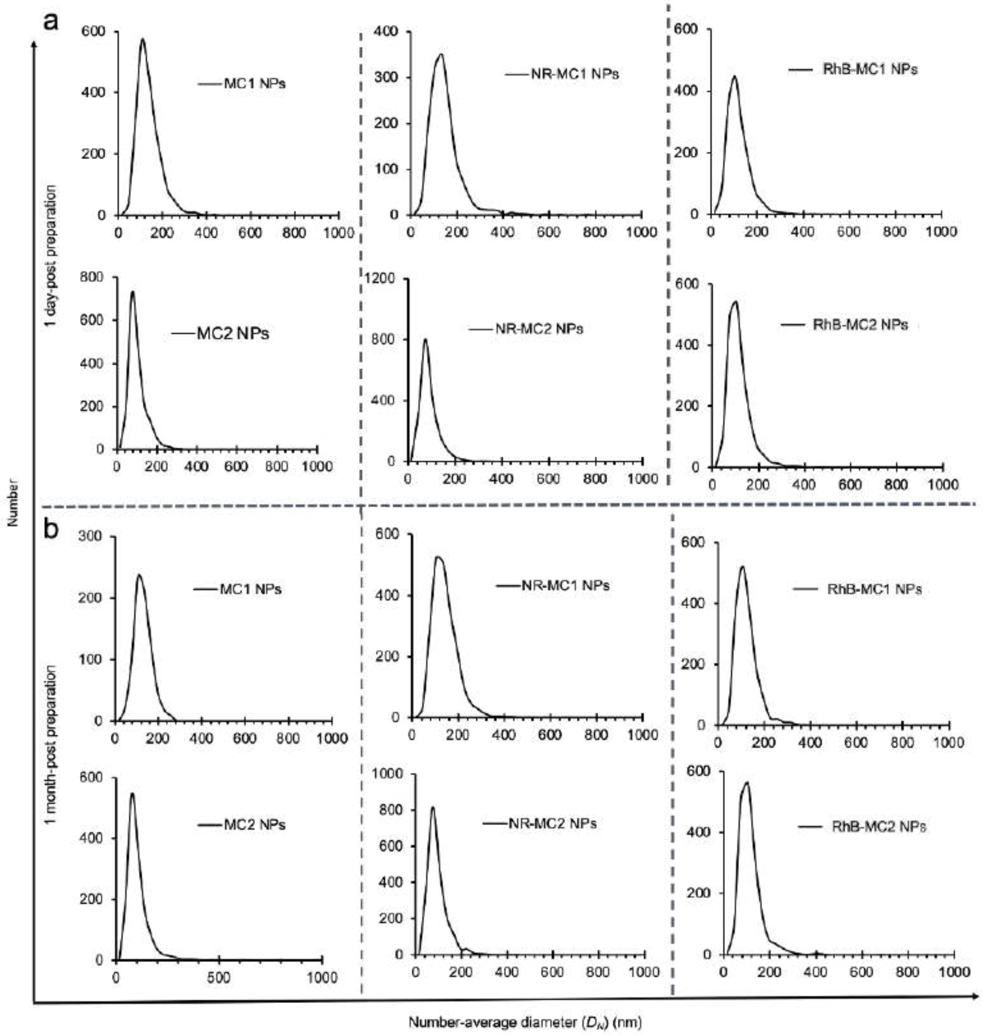

Figure S2. In vitro MTS assay of different concentrations (i.e. up to 3000 μg/mL) of unloaded MC1 and MC2 NPs incubated in adipose stem cells for 24 h displaying high cell viability.

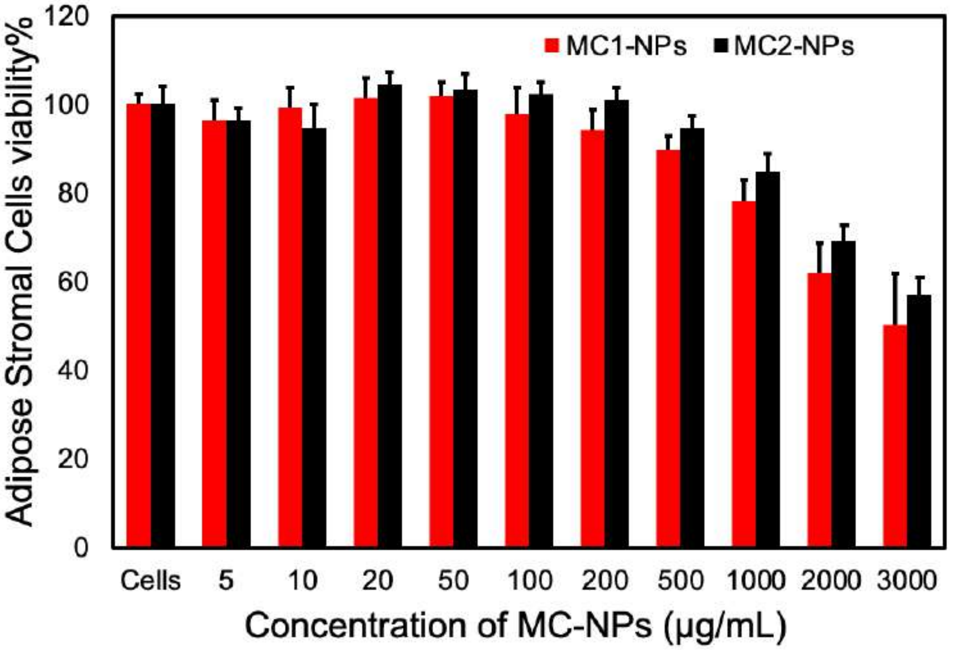

Figure S3. Cell viability studies (measured by CCK-8 assay) of RAW264.7 macrophage cells treated with various concentrations of blank MC NPs, Palb-MC NPs, and free Palb drug. The cells were exposed to MC NPs for 24 h.

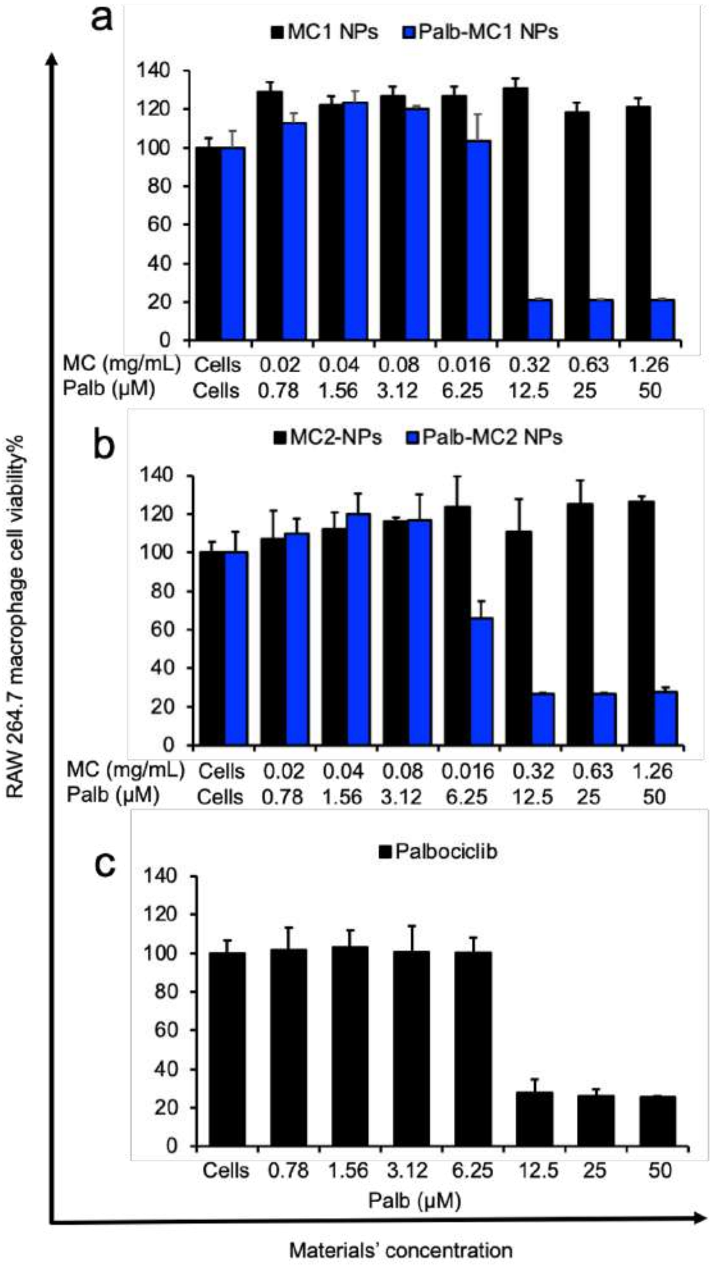

Table S2. IC50 values of MC toxicity curves against RAW264.7 cells calculated by GraphPad Prism, inhibitor vs. normalized response equation, the experiment was done in triplicate, and MC (mg/mL) and Palb (μM). It is noted that MC NPs themselves have not induced toxicity at higher concentrations so the IC50 is minimal (cannot be detected).

| | MC1 | Palb-MC1 | MC2 | Palb-MC2 | Free Palb |
|---|---|---|---|---|---|
| 24 h Treatment | -- | Palb = 16.64<br>MC1 = 0.42 | -- | Palb = 14.20<br>MC2 = 0.36 | 15.60 |
| 72 h Treatment | -- | Palb = 3.09<br>MC1 = 0.08 | -- | Palb = 2.81<br>MC2 = 0.07 | 8.00 |

Figure S4. (top) Fluorescence imaging studies were performed on model U-87 MG glioblastoma cells after treatment with MC1 and MC2 NPs. (A, B) controls, MC NPs and free NR dye, respectively. (C, D) NR-MC1 NPs at 4 h and 24 h incubation time, respectively. (E, F) NR-MC2 NPs at 4 h and 24 h incubation time, respectively. Scale bars are 100 and 200 μm in length as indicated in all inlets. (bottom) Flow cytometry data represent cellular uptake quantifications of NR-MC NPs in U-87 MG cells as compared to positive and negative controls.

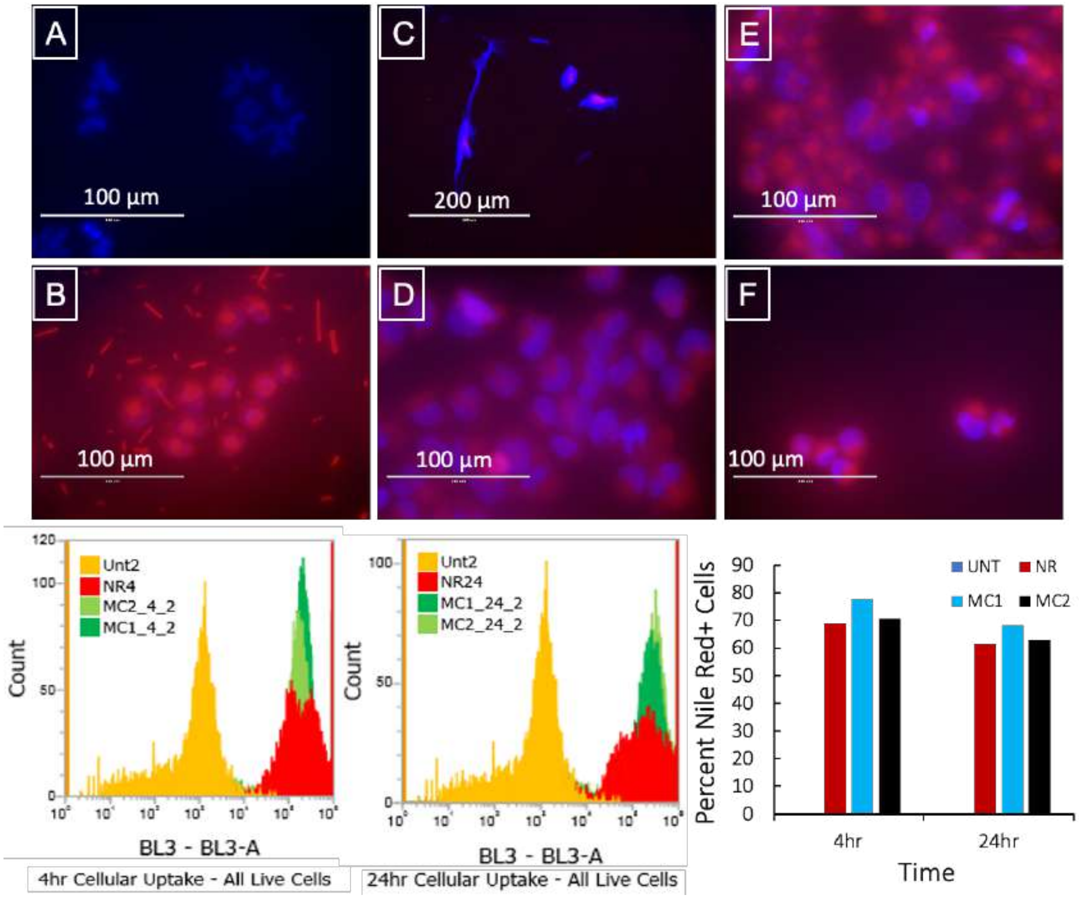

Figure S5. Cell viability studies (measured by CCK-8 assay) of HDFn cells treated with various concentrations of blank MC NPs. The cells were exposed to MC NPs for 24 h. The experiment was done in triplicates.

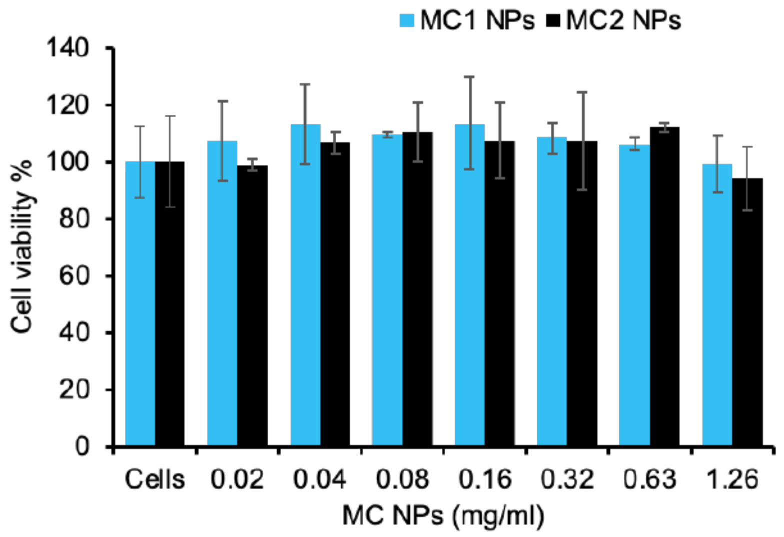


Table S3: IC50 values of MC toxicity curves against M14, Mel 29.1, and U-87 MG cells calculated by GraphPad Prism, inhibitor vs. normalized response equation, The experiment was done in triplicates, and the concentration of MC is mg/mL.

| | M14 cells | | Mel 29.1 cells | | U-87 MG cells | |
|---|---|---|---|---|---|---|
| | MC1 | MC2 | MC1 | MC2 | MC1 | MC2 |
| 72 h Treatment | 0.22 | 0.43 | 0.09 | 0.16 | 0.26 | 1.01 |
